# A Bayesian Model-Selection Approach for Determining the Number of Spectral Peaks in Neural Power Spectra

**DOI:** 10.1101/2024.08.01.606216

**Authors:** Luc E. Wilson, Jason da Silva Castanheira, Benjamin Lévesque Kinder, Sylvain Baillet

## Abstract

Neurophysiological brain activity comprises rhythmic (periodic) and arrhythmic (aperiodic) signal elements, which are increasingly studied in relation to behavioral traits and clinical symptoms. Current methods for spectral parameterization of neural recordings rely on user-dependent parameter selection, which includes selecting the correct maximum number of spectral peaks to fit to the spectrum. This challenges the replicability and robustness of findings. Here, we introduce a data-driven model-selection procedure for determining the appropriate number of oscillatory peaks to fit to neural power spectra, based on the Bayesian Information Criterion (BIC). We present extensive tests of the approach with ground-truth and empirical magnetoencephalography recordings. Data-driven model selection enhances both the specificity and sensitivity of spectral decompositions. Overall, the proposed spectral decomposition with data-driven model selection reduces reliance on user-defined ‘maximum number of peaks’ settings, enabling more robust, reproducible, and interpretable spectral parameterizations.

**Lay summary:** Brain activity is composed of rhythmic patterns that repeat over time and arrhythmic elements that are less structured. Recent advances in brain signal analysis have improved our ability to distinguish between these two types of components, enhancing our understanding of brain signals. However, current methods require users to adjust several parameters manually to obtain their results. The outcomes of the analyses, therefore, depend on each user’s decisions and expertise. To improve the replicability of research findings, the authors propose a revised method to streamline the analysis of brain signal contents. They developed a new algorithm that defines the parameters of the analytical pipeline informed by the data. The effectiveness of this method is demonstrated with both synthesized and real-world data. The approach is made available to all researchers as a free, open-source app, observing best practices for neuroscience research.

## 1 Introduction

Neural oscillations are rhythmic (periodic) signal components ubiquitously observed in electrophysiology across spatial and temporal scales (Buzsaki & Watson, 2012). In the power spectrum, periodic components can be modelled as Gaussian-shaped peaks emerging from an arhythmic (aperiodic) background (Wen & Liu, 2016; Donoghue et al., 2020; Wilson et al., 2022). Aperiodic activity is spectrally characterized by a reciprocal distribution of signal power that decays with frequency according to a power law (1/f^α^). In practice, the scalar exponent parameter α and broadband offset of the aperiodic model are inferred from estimates of the signal’s power spectrum density (PSD) of the electrophysiological signal. Computational neuroscience models and growing empirical evidence suggest that α reflects the physiological balance between excitatory (E) and inhibitory (I) neural activity (Brake et al., 2024; Chini et al., 2022; Gao et al., 2017; Wiest et al., 2023), and the offset is related to aggregate neuronal population spiking (Miller et al., 2014; Voytek & Knight, 2015). These model parameters of the aperiodic spectral component decrease with age, accounting for the observation of a *flatter* power spectrum in aging (Cellier et al., 2021; Donoghue et al., 2020; Voytek et al., 2015). They also fluctuate during cognitive tasks (Donoghue et al., 2020; Gyurkovics et al., 2022; Preston et al., 2022; Waschke et al., 2021) and reflect behavioral traits (Ostlund et al., 2021; Wilson et al., 2022).

Recent algorithms and software such as *specparam* (Donoghue et al., 2020) and Spectral Parameterization Resolved in Time (*SPRiNT;* Wilson et al., 2022) have streamlined the adoption of spectral parameterization in electrophysiological research. These tools require users to define a number of method parameters (hyperparameters), such as model complexity via the pre-specification of the maximum number of spectral peaks N_G_ to be adjusted from the empirical PSD (Gerster et al., 2022; Ostlund et al., 2022; Wilson et al., 2022). When hyperparameters are not set appropriately, the *specparam* algorithm either fits spurious, outlier spectral peaks or misses genuine spectral peaks (Donoghue et al., 2020). Similarly, time-resolved spectral parametrization tools rely on hyperparameters to minimize the detection of outlier spectral peaks (Brady & Bardouille, 2022; Kosciessa et al., 2020; Seymour et al., 2022; Stokes et al., 2023; Whitten et al., 2011; Wilson et al., 2022).

Setting model hyperparameters is a prevalent challenge across many fields of science and engineering. Good-practice approaches recommend prioritizing parsimonious models with a balance between simplicity (fewer hyperparameters) and the ability to fit the observed data (more flexibility; Vandekerckhove et al., 2015). Limiting the number of spurious peaks fit to the spectrum is crucial, as every additional peak fit to the spectrum increases the number of model parameters by three (i.e., peak center frequency, amplitude, and width), which greatly increases the complexity of the model.

To address this concern, we propose a principled model selection strategy for the parameterization of neural power spectra. The method proceeds with adjusting progressively the number of Gaussian peaks fit to the empirical data spectrum (i.e., the model complexity) and determines the parameters of the simplest model that adequately accounts for the data using the Bayesian information criterion (BIC). This approach allows us to determine the simplest model that adequately explains the data, without requiring users to set a maximum number of peaks a priori. This also enables the quantitation of evidence for periodic activity in spectral data via Bayes factor analysis. We demonstrate below the method’s performance using extensive ground-truth simulated data and a large set of empirical resting-state magnetoencephalography (MEG) from N=606 participants.

## 2 Methods

### 2.1 Model Selection using Bayesian Information Criterion

In the context of parsimonious modeling of power spectra and spectrograms, our goal is to optimize the trade-off between model fidelity and complexity. This principle emphasizes deriving the most accurate and representative model directly from empirical data while minimizing the inclusion of unnecessary assumptions or parameters (Myung, 2000). Among the methods for comparing models, Bayes factors are noteworthy as they balance model fit evaluation with the principle of simplicity (Jefferys & Berger, 1992). These factors can be effectively estimated using the Bayesian Information Criterion (BIC), which offers a pragmatic trade-off between goodness-of-fit and complexity in terms of the number of model parameters (Schwarz, 1978; Vandekerckhove et al., 2015).

The primary motivation is to select the model with the most parsimonious fit: as such, we opted against Akaike Information Criterion (AIC), which tends to favour more complex models. We similarly did not explore other goodness-of-fit metrics, as BIC affords us the ability to easily compute Bayes Factors evidence for periodic activity in the power spectrum. Note, the outputs of the algorithm include the log-likelihoods of each model tested, which allows users to convert BIC metrics to AIC if desired.

The primary innovation of ms-specparam is a new objective function that is being minimized. In the *specparam* approach, model fitting involves minimizing the least-squares error between model predictions and the empirical power spectrum. The *ms-specparam* method refines this objective by estimating the negative log-likelihood of a model:

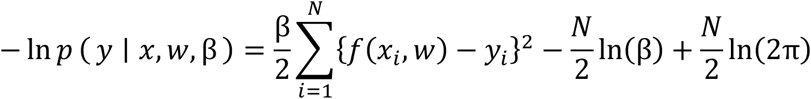

where (*x*) and (*y*) are the frequency bins and empirical spectral power values, respectively; (*f*(*x*_*i*_, *w*)) is the spectral power predicted by the model at frequency (*x*_*i*_); (*y*_*i*_) is the empirical spectral power value at frequency (*x*_*i*_); (*w*) represents the model parameters, (*N*) the number of frequency bins. (β) is the precision, or inverse variance, of residuals.

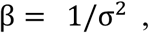

Under the assumption of zero-mean (unbiased) Gaussian noise in the empirical power spectrum, minimizing negative log-likelihood provides an equivalent solution to minimizing squared error (Mitchell, 1997). Finally, we express the *ms-specparam* optimization’s output in terms of the Bayesian information criterion (BIC):

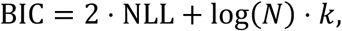

where NLL is the negative log-likelihood, (*N*) is the number of frequency bins, and (*k*) represents the total number of parameters, which includes the aperiodic parameters (exponent and offset) and three additional parameters for each peak (center frequency, amplitude, bandwidth). Note that (*k* = 3*P* + 2), where (*P*) represents the number of peaks.

The *ms-specparam* approach, much like the *specparam* algorithm, iteratively fits models of increasing complexity by adding peaks and minimizing the squared error. Note that for each additional modelled Gaussian peak, the number of parameters increases by three (i.e., the peak center frequency, width, and amplitude). As in the default parameterization procedure, peaks are fit in decreasing order of amplitude, where an additional peak is fit if it satisfies criteria for amplitude (i.e., ‘minimum peak height’ and ‘relative peak threshold’) as well as distance from other peaks or the spectrum edges. However, *ms-specparam* performs an additional, combined bounded optimization of both aperiodic and periodic parameters, and converts the squared error into a negative log-likelihood, which is then used to calculate the BIC. To identify the model with the lowest BIC, ms-specparam saves optimized models of the spectrum following the addition of each spectral peak, as well as their respective BIC. Finally, *ms-specparam* compares and provides the parameters of the model with the lowest BIC, thus achieving a balance between fit quality and model simplicity. As such, our approach optimizes the set of peaks detected through the original *specparam* algorithm in order of decreasing peak amplitude, based on BIC. In fact, the result is nearly equivalent algorithmically to individually setting the ‘maximum number of peaks’ hyperparameter in *specparam* for each spectrum to the quantity determined by *ms-specparam* to provide the lowest BIC, with the benefit of being data-driven and automatic.

We share *ms-specparam* as a MATLAB plug-in library that interoperates with *Brainstorm* (Tadel et al., 2011), as well as an equivalent Python library. Both implementations are open-source and accessible to all users.

### 2.2 Algorithm Settings and Hyperparameters

The spectral parameterization of neural power spectra (synthetic and empirical) was conducted in the frequency range of 1-40 Hz using three distinct approaches, each characterized by different hyperparameter settings:

#### Default Hyperparameters

This approach adhered to the default settings established in the Python implementation of *specparam* (Donoghue et al., 2020), with an increased range for peak width limits. The hyperparameters included a minimum peak height of 0.1 arbitrary units (a.u.), a maximum of 6 peaks, peak width limits set within the range of [1, 24] Hz, and a gaussian overlap threshold of 0.75 s.d.

#### ms-specparam

This approach used the same hyperparameter settings as *default-specparam*, including a minimum peak height of 0.1 a.u., a maximum of 6 peaks, peak width limits between [1, 24] Hz, and a gaussian overlap threshold of 0.75 s.d. The most parsimonious spectral model is selected according to the procedure described in *Model Selection using Bayesian Information Criterion*.

### 2.3 Synthetic Data

We created 5,000 synthetic neural power spectra using a range of aperiodic parameters: exponents from 0.5 to 2 Hz^-1^) and offsets from −8.1 to −1.5 arbitrary units (a.u.). Each power spectrum was augmented with zero to four peaks, with 1,000 instances for each peak quantity. The parameters for these peaks fell within specified ranges: center frequencies between 3 and 35 Hz, amplitudes from 0.1 to 1.5 a.u., and bandwidths (2 s.d.) from 2 to 6 Hz. We ensured a minimum separation of one bandwidth between adjacent peak frequencies. The frequency domain for simulation spanned from 0.5 to 100 Hz with increments of 0.5 Hz.

To mimic realistic noise conditions, we introduced Gaussian white noise at varying intensities: low (0.05 a.u.), medium (0.10 a.u.), and high (0.15 a.u.). We then applied *default-specparam* (minimum peak height of 0.1 a.u., up to six peaks, peak width range of 1 to 24 Hz, and gaussian overlap threshold of 0.75 s.d.) alongside *ms-specparam* with identical parameters.

For accuracy assessment, a ‘hit’ was designated when an identified peak’s center frequency was within one bandwidth (2 s.d.) of a true peak. In cases where multiple identified peaks met this criterion, the peak with the highest amplitude was selected as the ‘hit.’ Peaks detected by the algorithm that did not correspond to a true peak were classified as false positives.

We defined peak sensitivity as the ratio of ‘hits’ to the total number of simulated peaks, and the positive predictive value as the ratio of ‘hits’ to all detected peaks (including false positives). For peaks identified as ‘hits,’ we calculated the parameter estimation error as the absolute deviation from the true values.

### 2.4 Bayes factor evidence for periodic brain activity

Bayes factor (BF) provides a statistical measure for comparing two models, offering evidence about the presence of periodic activity within neural power spectra. The computation of the Bayes factor follows the formula proposed by Wagenmakers (2007):

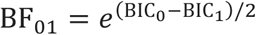

where (BIC_0_) is the Bayesian Information Criterion (BIC) value for the aperiodic-only model and (BIC_1_) is the BIC value of the lowest-BIC model.

In this context, (BIC_0_) represents the BIC for the aperiodic-only model, which posits that the data can be explained without invoking rhythmic components in the power spectrum. (BIC_1_), on the other hand, corresponds to the model that includes both aperiodic and periodic elements and has the lowest BIC among all models considered.

The Bayes factor (BF_01_) compares these models, translating the difference in their BIC values into the odds ratio against periodic activity in the power spectrum. A smaller (BF_01_) implies stronger evidence against the aperiodic-only model, thereby indicating the presence of significant periodic activity within the brain’s neural power spectra. Conversely, larger (BF_01_)values suggest that the periodic components do not significantly improve the model beyond the aperiodic activity alone.

This approach allows for the quantitative assessment of oscillations in neural recordings, providing a more rigorous foundation for claims of rhythmic brain activity observed in electrophysiological data.

### 2.5 Empirical data

The empirical dataset for our study was obtained from the Cambridge Centre for Aging Neuroscience repository (Cam-CAN; Shafto et al., 2014; Taylor et al., 2017). This comprehensive dataset includes 606 healthy individuals aged 18 to 90 years (mean age = 54.69; SD = 18.28), with a balanced gender representation (299 females). Each participant underwent a thorough assessment, beginning with a detailed home interview followed by a resting-state magnetoencephalography (MEG) session. The MEG recordings, lasting approximately 8 minutes each, were conducted using a 306-channel VectorView MEG system (MEGIN). These recordings were complemented with structural T1-weighted magnetic resonance imaging (MRI) to provide anatomical context for MEG source mapping. All data collection occurred at a single, consistent location to maintain uniformity in data acquisition.

This rich dataset forms the basis for our analyses, allowing for a comprehensive investigation into the spectral properties of neural signals across a wide age spectrum.

#### 2.5.1 MEG Preprocessing and Source Mapping

We preprocessed magnetoencephalography (MEG) data with Brainstorm (Tadel et al., 2011; March 2021 distribution), integrated with MATLAB (2020b; Natick, MA), adhering to established best-practice guidelines (Gross et al., 2015). The preprocessing methodology followed protocols detailed previously (da Silva Castanheira et al., 2024a; da Silva Castanheira et al., 2024b).

Line noise artifacts at 50 Hz and its first 10 harmonics were filtered using a notch filter bank. Additionally, an 88-Hz artifact present in the Cam-CAN dataset (Wiesman et al., 2022) was removed. To address slow-wave and DC-offset artifacts, a high-pass finite impulse response (FIR) filter with a cutoff frequency of 0.3 Hz was applied. Signal-Space Projections (SSPs) were implemented to attenuate cardiac artifacts and mitigate low-frequency (1–7 Hz) and high-frequency (40–400 Hz) noise components, typically originating from saccades and muscle activities.

Brain source models were anchored to the individual T1-weighted MRI data of each participant. Automatic segmentation and labeling of MRI volumes were achieved using FreeSurfer (Fischl, 2012). Co-registration with MEG data was facilitated using approximately 100 head points digitized for each participant. MEG biophysical head models were computed using *Brainstorm*’s overlapping-spheres model (default parameters).

Cortical source models were estimated using linearly constrained minimum-variance (LCMV) beamforming, following *Brainstorm*’s default parameters (2018 version for source estimation processes). MEG source orientations were constrained normal to the cortical surface, distributed across 15,000 locations. Neural power spectra were then calculated for each of the 148 cortical regions defined by the Destrieux atlas (Destrieux et al., 2010). These calculations were based on the first principal component of all signals within each region of interest (ROI). Neural spectral power was estimated using Welch’s method, utilizing 2-second windows with a 50% overlap. We parameterized the resulting neural power spectra with *default-specparam* (minimum peak height of 0.1 a.u., up to six peaks, peak width range of 1 to 24 Hz, and gaussian overlap threshold of 0.75 s.d.) alongside *ms-specparam* with identical parameters. We chose these hyperparameters based on visual inspection of the data.

#### 2.5.2 Statistical Analyses

To evaluate the accuracy of spectral parameters generated by *ms-specparam*, we employed two-sample non-parametric permutation t-tests from the *RVAideMemoire* package in R. This statistical approach allowed us to test differences in spectral parameter estimates, specifically focusing on five key variables: aperiodic exponent, aperiodic offset, peak center-frequency, peak amplitude, and peak bandwidth.

To capture differences in residual variance and the number of peaks fitted between algorithms in our empirical data, we similarly relied on paired non-parametric permutation t-tests.

We implemented hierarchical linear regression models using the *lmer* function in R to test how algorithm choice impacts age’s effect on both aperiodic exponent and offset. The models were formulated as:

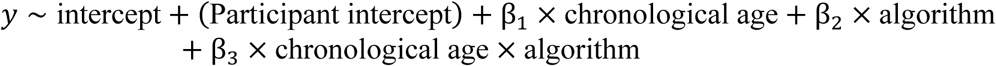

where (*y*) represents the dependent variables, including the aperiodic exponent and offset.

Separate linear regression analyses were conducted to compare *ms-specparam* against *default-specparam* for each dependent variable of interest. Chronological age and the chosen algorithm (e.g., *ms-specparam* vs. *default-specparam*) were introduced as fixed predictors, and individual participants were treated as a random factor to account for inter-individual variability. Note that we report standardized beta coefficients which can be interpreted as effect sizes, and otherwise report effect sizes using Cohen’s D.

## 3 Results

Figure 1a highlights how user-dependent hyperparameters affect the outcome of spectral parameterization. In the example shown, the hyperparameter specifying the maximum number of peaks expected from spectral parameterization was set manually to a value of 6. However, the spectrum of the present empirical data contains only two peaks (left panel; Figure 1a). Under high signal-to-noise conditions, spectral parameterization may yield only two peaks as expected. However, when data is realistically noisy, the spectral parameterization algorithm may overestimate the number of peaks in the spectrum to account for noise-related fluctuations (right panel; Figure 1a). Ideally, spectral and spectrogram parameterization should automatically and adaptively adjust model hyperparameters to account for the noise level in the data.

**Figure 1:**
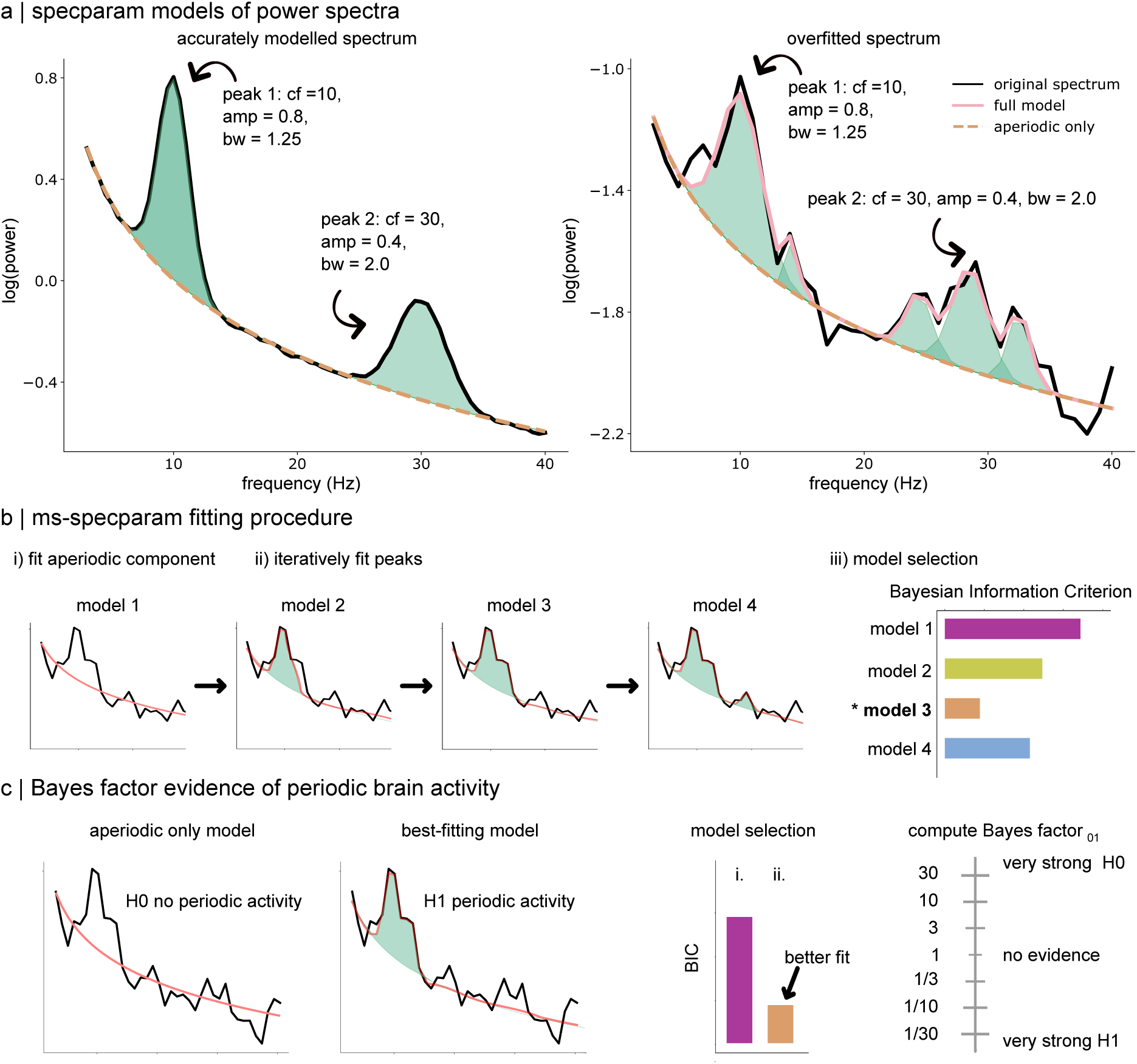
Spectral Parameterization with Model Selection. (a) Illustration of a spectral parameterization of a simulated power spectral density estimate (black line) obtained with *specparam* in the context of lower (left panel) and higher (right panel) noise levels. Both spectra are generated using the same spectral parameters (i.e., two spectral peaks). In more noisy conditions, *specparam* (pink line) fits a greater number of spectral peaks (green shaded areas) than what is present (simulated) in the data, resulting in overfitting (right panel). Key: ‘*cf’* refers to a peak’s center frequency, ‘*amp*’ refers to a peak’s amplitude, and ‘*bw*’ refers to a peak’s bandwidth. (b) *ms-specparam* is a method for spectral parameterization combined with a model selection procedure. It first adjusts a model for the aperiodic component of the spectrum (subpanel i) before adding spectral peaks (green shaded areas) in an iterative fashion (subpanel ii). These successive models are then assessed via the Bayesian Information Criterion (BIC; subpanel iii). (c) The resulting BIC model is then subjected to Bayes factor inference against the aperiodic spectral model (panel i) to adjudicate whether spectral peaks are likely to be present in the data power spectrum. A Bayes factor greater than 1 indicates evidence in favour of periodic components in the power spectrum over the aperiodic-only model (panel iv).

To address these issues, we propose an approach that proceeds iteratively with the parameterization of empirical power spectra and spectrograms with increasing model complexity (i.e., the maximum number of spectral peaks). We derive the Bayesian Information Criterion (BIC) for each hyperparameter setting. The most parsimonious hyperparameter setting to model the empirical power spectrum is the one corresponding to the lowest BIC value (Figure 1b). If the most parsimonious model contains spectral peaks, we can further quantify the evidence for periodic activity using the Bayes factor relative to the aperiodic-only model (Figure 1c). Our method, coined *ms-specparam*, is freely available through *Brainstorm* (Tadel et al., 2011) and on GitHub (github.com/lucwilson/model_selection).

We tested and validated *ms-specparam* using 5,000 ground-truth, synthetic but neurophysiologically plausible power spectra. We compared its performance against the original *specparam* algorithm, configured to its default hyperparameters (referred to as *default-specparam*; see Methods). We also applied *ms-specparam* to task-free MEG recordings from 606 participants to replicate, with less dependence on user-selected hyperparameters, the previously reported findings of an age-related decline in the aperiodic exponent of the neurophysiological power spectrum (Voytek et al., 2015).

### 3.1 Synthetic, Ground-Truth Data

Each of the 5,000 synthetic power spectra comprised an aperiodic component, with offset values ranging from −8.1 to −1.5 arbitrary units (a.u.) and exponents set between 0.5 and 2 Hz^-1^. We randomly added between 0 and 4 spectral peaks to the aperiodic background of each power spectrum (1,000 simulated spectra for each number of peaks; see Methods). Zero-mean Gaussian noise was added to the resulting power spectra with varying standard deviation values (s.d.).

In moderate noise conditions (s.d. = 0.10), *ms-specparam* demonstrated a slightly lower sensitivity (89%) in detecting spectral peaks compared to *default-specparam* (91%). However, it had a substantially higher positive predictive value for peak detection (*ms-specparam*: 96%; *default-specparam*: 63%). On average, *default-specparam* overestimated the number of peaks in the spectrum by 59%, whereas *ms-specparam* underestimated the number of peaks by 13% (Figure 2a). We observed similar results for sensitivity and PPV in both algorithms at lower and higher noise levels (Figures S1, S2).

**Figure 2:**
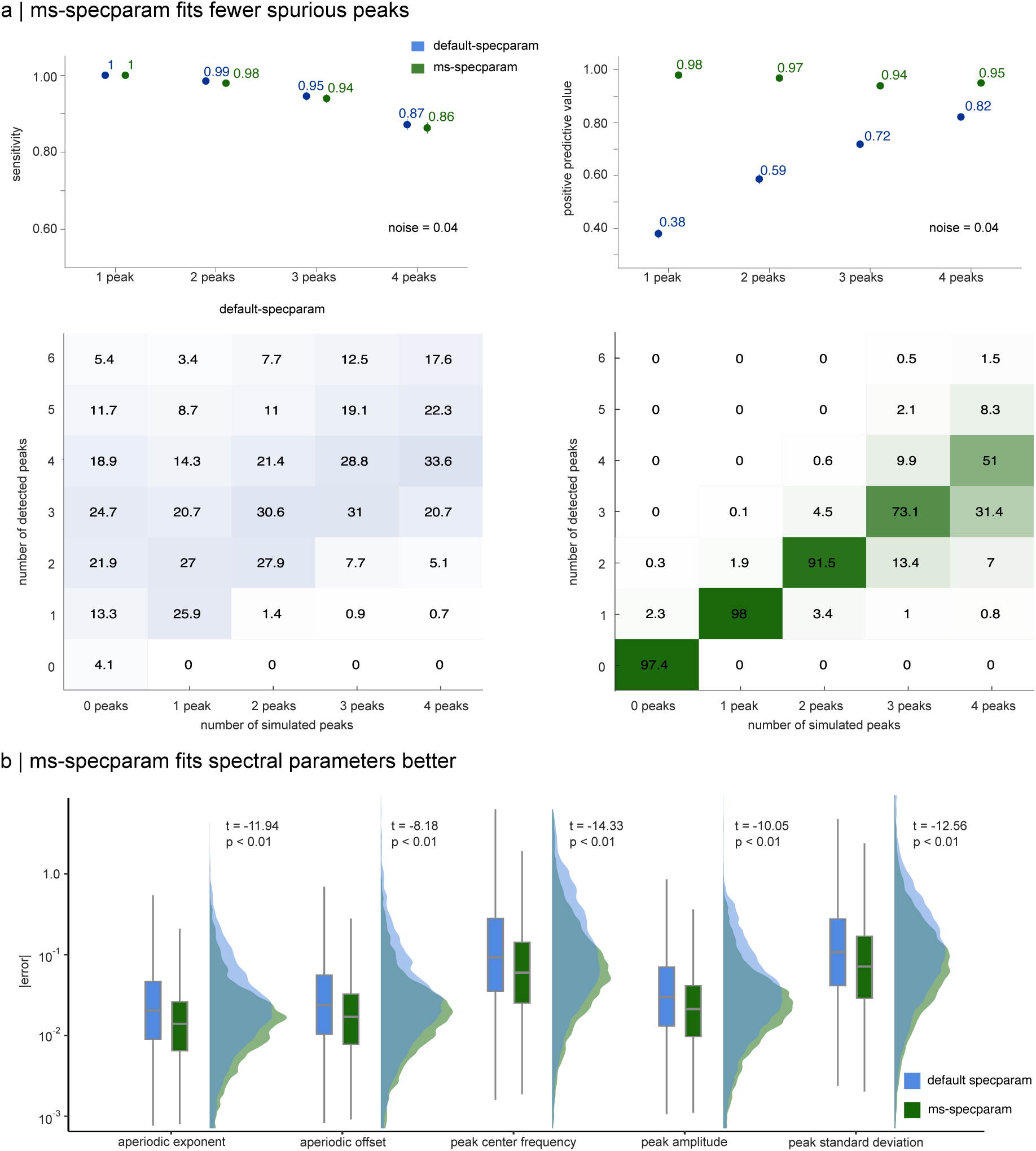
Performances on Synthetic Stationary Data. (a) Sensitivity and positive predictive value (PPV) for detection of spectral peaks (top row). *ms-specparam* (green) has similar sensitivity (89%) than *default-specparam* (blue; 91%), but superior PPV (96% vs. 63%). The heat maps below report the ground-truth vs. estimated number of spectral peaks (with percent incidence listed in each element) and highlight *ms-specparam*’s improved peak detection accuracy. (b) Boxplots and empirical density distributions reporting the errors on the estimates of the spectral parameters derived using *ms-specparam* and *default-specparam*. For every spectral parameter, *ms-specparam* estimated values with significantly lower mean absolute error (one-tailed permutation t-test, all p<0.05).

We found that, on average, all parameter estimates derived from *ms-specparam* are more accurate than those derived from *default-specparam*: mean error in aperiodic exponent (t = - 11.94, p = 0.0019, Cohen’s D = −0.24; one-tailed permutation t-tests), aperiodic offset (t = −8.18, p = 0.0019, Cohen’s D = −0.16), peak center frequency (t = −14.33, p = 0.0019, Cohen’s D = - 0.21), peak amplitude (t = −10.05, p = 0.0019, Cohen’s D = −0.15), and peak bandwidth (t = - 12.56, p = 0.0019, Cohen’s D = −0.18; Figure 2b).

We then sought to characterize which spectral peak features may impact the detection of spectral peaks. First, we explored whether the distance between two spectral peaks negatively impacts their detection. To do so, we computed the maximum distance between two spectral peaks within a simulation, and assigned it to a quantile. We observed that for synthetic data containing 2 simulated spectral peaks, ms*-specparam* demonstrated similar sensitivity (92%) in detecting spectral peaks compared to *default-specparam* (96%) for moderate noise conditions (SNR = 0.04; Figure 3a). Distance between spectral peaks (i.e., quantiles) did not significantly impact sensitivity across all simulations. Similarly, we observed minimal effects of distance between spectral peaks on PPV (Figure 3b).

**Figure 3:**
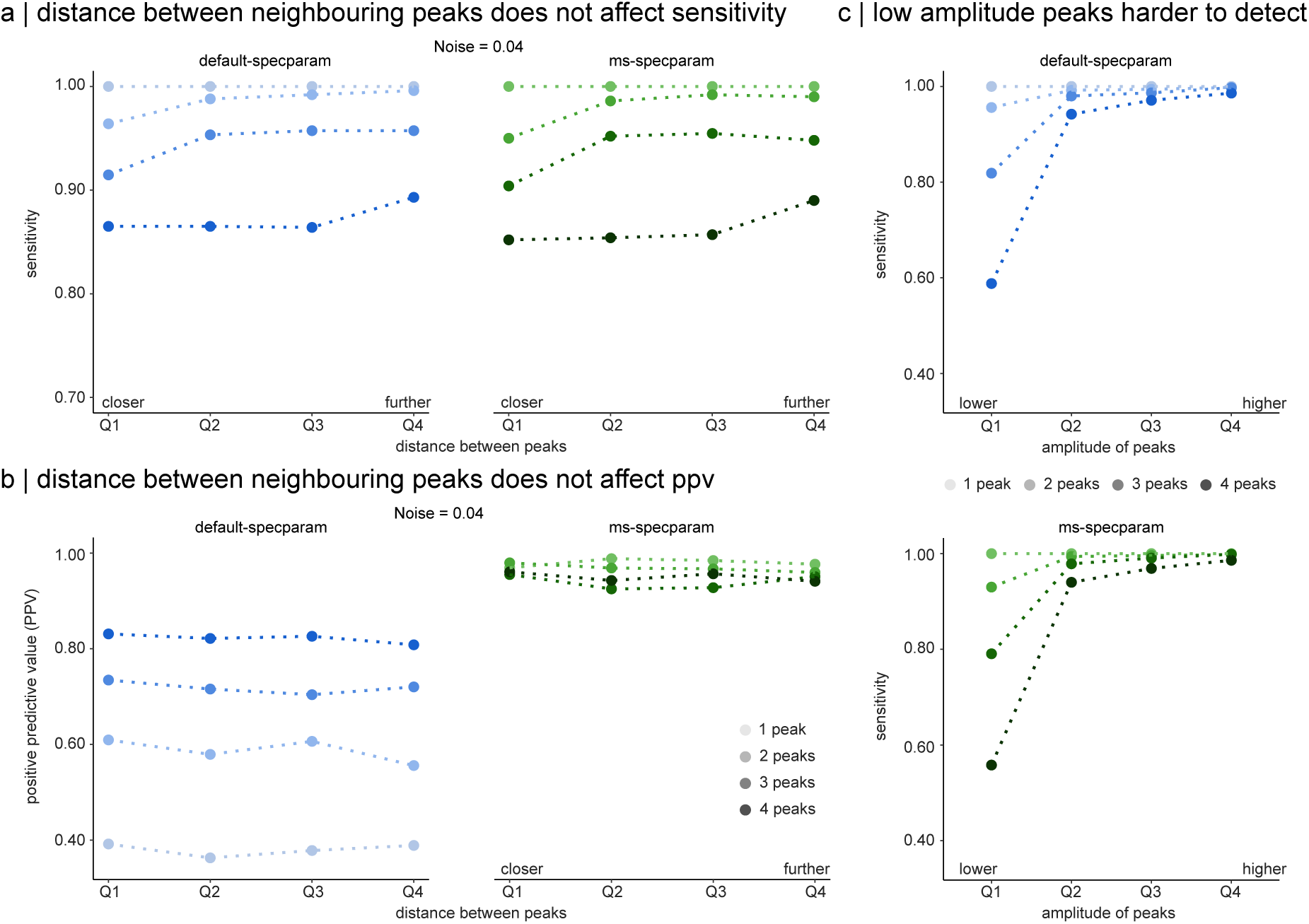
Larger Spectral Peak Amplitude Increases Peak Detection. (a) Sensitivity for detection of spectral peaks for increasing distances between synthetic spectral peaks (quartiles; x-axis). *ms-specparam* (green) has a similar sensitivity to *default-specparam* (blue) regardless of the distance between two spectral peaks (x-axis). Saturation of the colours indicates the number of synthetic spectral peaks. (b) Positive predictive value (PPV) for detection of spectral peaks for increasing distances between synthetic spectral peaks (quartiles; x-axis). *ms-specparam* (green) outperforms *default-specparam* (blue) for all quartiles of distances between peaks. (c) Sensitivity for detecting spectral peaks for the quartiles of peak amplitude (x-axis). Both *default-specparam* and *ms-specparam* show reduced sensitivity for detecting spectral peaks when they are of low amplitude. This effects if more pronounced for simulations with multiple spectral peaks (dark coloured points).

Second, we explore the impact of spectral peak amplitude on spectral peak detection. We similarly computed sensitivity for each quantile of peak amplitude from lowest amplitude peaks (Q1) to the highest amplitude peaks (Q4). We observed that both algorithms showed reduced sensitivity to spectral peaks for low amplitude oscillations, with both algorithms demonstrating lowest sensitivity for spectral peak detection for synthetic data with four spectral peaks (Figure 3c). We note that the effect of amplitude on peak detection became increasingly pronounced with additional synthetic spectral peaks (darker lines, Figure 3c).

### 3.2 Empirical MEG Data

We applied *ms-specparam* to resting-state MEG data from the Cam-CAN repository (N=606; Shafto et al., 2014; Taylor et al., 2017). We first preprocessed and source-mapped the MEG time series using *Brainstorm* (Tadel et al., 2011) following good-practice guidelines (Gross et al., 2013). We then derived the PSDs of each cortical parcel of the Destrieux atlas (Destrieux, 2010; see Methods). We then compared models generated with *ms-specparam* to those from *specparam*, using hyperparameter settings: *default-specparam* (minimum peak height: 0.1 a.u.; maximum number of peaks: 6; peak width limits: [1 24]; gaussian overlap threshold: 0.75 s.d.).

We found that *ms-specparam* generated models with less residual variance (i.e., mean squared error, MSE; average MSE = 1.33×10^-3^, SD = 4.42×10^-4^) than *specparam* settings (default: average MSE = 2.82×10^-3^, SD = 1.01×10^-3^; t = −48.11, p < 0.001, Cohen’s D = −1.90; Figure 4a). This observation was consistent across all cortical parcels, with posterior parietal areas showing the greatest enhancements in model goodness-of-fit (Figure 4a). Reduced residual variance was consistent over the entire frequency range (1–40 Hz), with marked improvements with respect to both *specparam* settings over the edges of the spectrum (<5 Hz and >35 Hz; Figure 4a). This held for spectral models fit to different frequency ranges (i.e., 1-45 Hz and 4-45 Hz; Figure S3).

**Figure 4:**
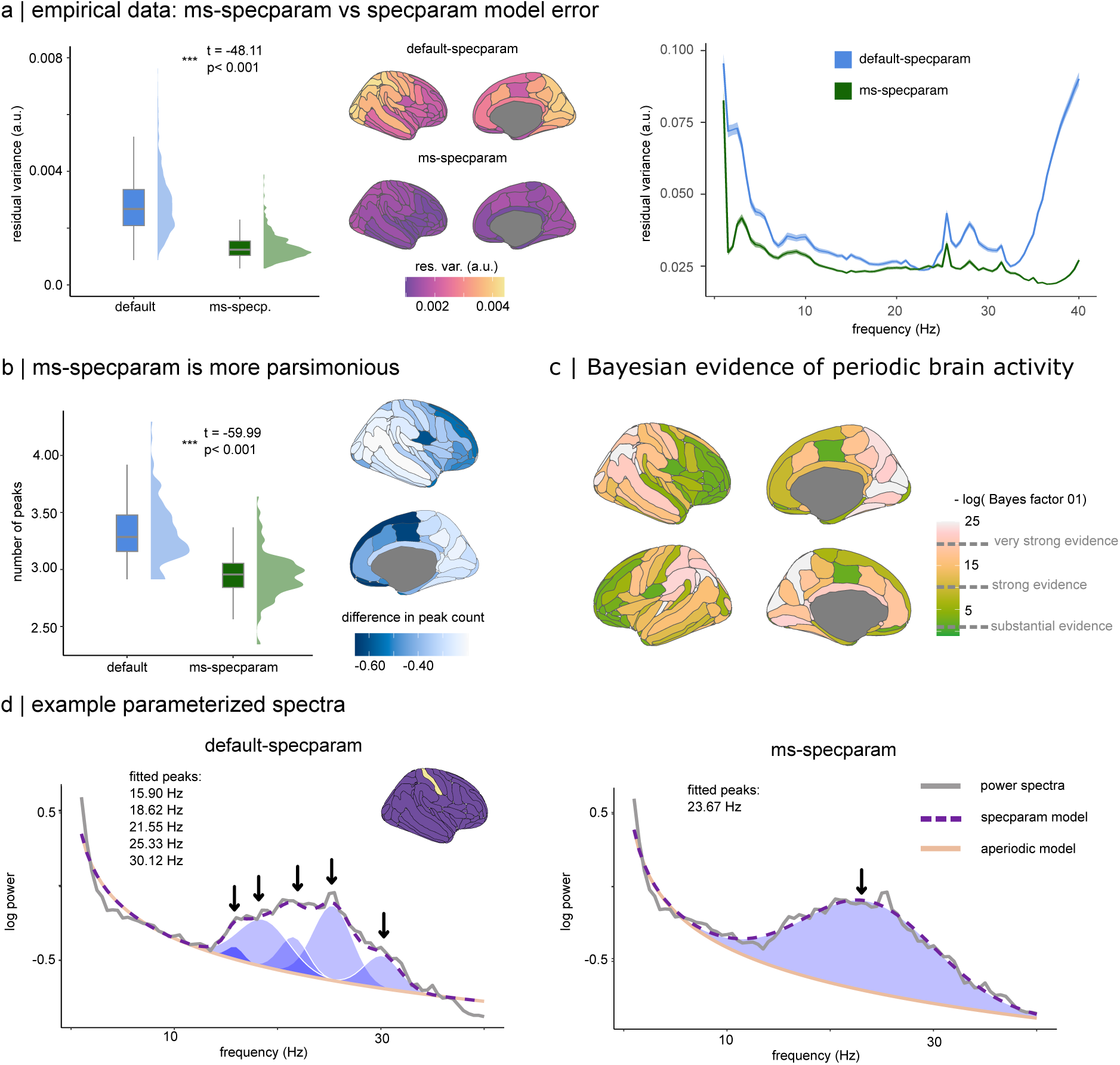
Performances on Empirical MEG Data. (a) Residual variance analysis across the 606 participants and all brain regions shows *ms-specparam* (green) with consistently lower residual variance, indicating a superior fit relative to default *specparam* (blue; left panel). The brain maps display residual variance values for each cortical parcel. A frequency breakdown (right panel) reveals *ms-specparam* outperforms default *specparam* variations across the spectrum, particularly at the edges of the frequency spectrum. (b) *ms-specparam* estimates less spectral peaks than *default-specparam*, demonstrating more parsimonious modelling, as reported in the box/density plots (left panel). The brain maps indicate that less spectral peaks (i.e., darker colours) were detected in posterior cortical parcels with *ms-specparam* (right panel). (c) Bayesian evidence for periodic, rhythmic brain activity mapped across the cortical surface emphasizes occipital and left temporal regions. (d) Parameterized spectra from the right post-central gyrus of a sample subject highlight the differences between algorithms: *ms-specparam* fits one peak (right panel), reflecting the dominant oscillations, whereas *default-specparam* fits five (left panel), some of which may be redundant or overfitted, as seen in the overlaid spectral models.

We also observed that *ms-specparam* detected, on average, −0.37 fewer spectral peaks than *default-specparam*, and therefore, as expected, provided more parsimonious spectral parameterizations (t= −59.99, p < 0.001, Cohen’s D = −0.81; Figure 4b). This was particularly evident in frontal brain regions, where *ms-specparam* fit 0.3 fewer peaks. We interpret these effects as default *specparam* fitting more lower amplitude spectral peaks that are common in frontal brain areas, particularly in the theta range (4-8 Hz) (Donoghue et al., 2020). In line with this interpretation, the spectral peaks fit by *ms-specparam* were of higher amplitude than *default*-*specparam* (default average: 0.36; ms-average 0.44; t= −120.56, p < 0.001, Cohen’s D = 0.32), suggesting greater selectivity.

With *ms-specparam*, we performed Bayes factor analyses (Vandekerckhove et al., 2015) as an objective measure of evidence for the presence of rhythmic activity in the neurophysiological power spectrum. We found that the bilateral cuneus exhibited the highest Bayesian evidence for rhythmic activity in the resting-state, while we found the lowest evidence of rhythmic activity in the medial frontal cortices (Figure 4c).

For illustration purposes, Figure 4d shows representative spectral parameterizations obtained from neurophysiological time series recorded the right post-central gyrus. In this instance, *ms-specparam* identified one spectral peak while *default-specparam* adjusted five spectral peaks (*default-specparam* MSE: 1.76×10^-3^; *ms-specparam:* 1.41×10^-3^). This example illustrates how *ms-specparam* reattributes variance previously modelled by several low amplitude oscillatory peaks back to the arrhythmic exponent to achieve a more parsimonious model.

With regard to periodic spectral parameters, we observed that *ms-specparam* fitted spectral peaks of larger bandwidths (*ms-specparam* average = 4.13, *default-specparam* average = 2.74; t = 192.28, p = 0.0019, Cohen’s D = 0.54) but of similar centre frequencies (*ms-specparam* average = 16.17, *default-specparam* average = 15.86; t = - 15.93, p = 0.0019, Cohen’s D = −0.04) in comparison to *default-specparam*.

### 3.3 Age-Related Flattening of the Aperiodic Spectrum Depends on Hyperparameter Choice

We aimed to replicate with *ms-specparam* previous observations using *specparam* of age-related decreases in aperiodic exponent (Donoghue et al., 2020; Voytek et al., 2015). We examined whether the choice of algorithm (i.e., *default-specparam* vs *ms-specparam*) influenced the detection of these aging effects. To do this, we fitted hierarchical linear regression models where age, the choice of spectral parameterization algorithm, and their interaction were included as predictors of both aperiodic exponent and aperiodic offset, respectively. A significant interaction would suggest that the relationship between chronological age and aperiodic exponent is weakened by the choice of parameterization algorithm.

We found that age-related decreases in aperiodic exponent are moderated by the algorithm used, whether *ms-specparam* or *specparam* (i.e., a significant interaction between age and spectral parameterization algorithm). This indicates that the observed age-related changes in aperiodic exponent may be influenced by the choice of parameterization method rather than solely reflecting genuine neurophysiological effects (default hyperparameters: β = 0.11, SE = 0.04, CI [0.04, 0.18]; Figure 5a and Table S1).

**Figure 5:**
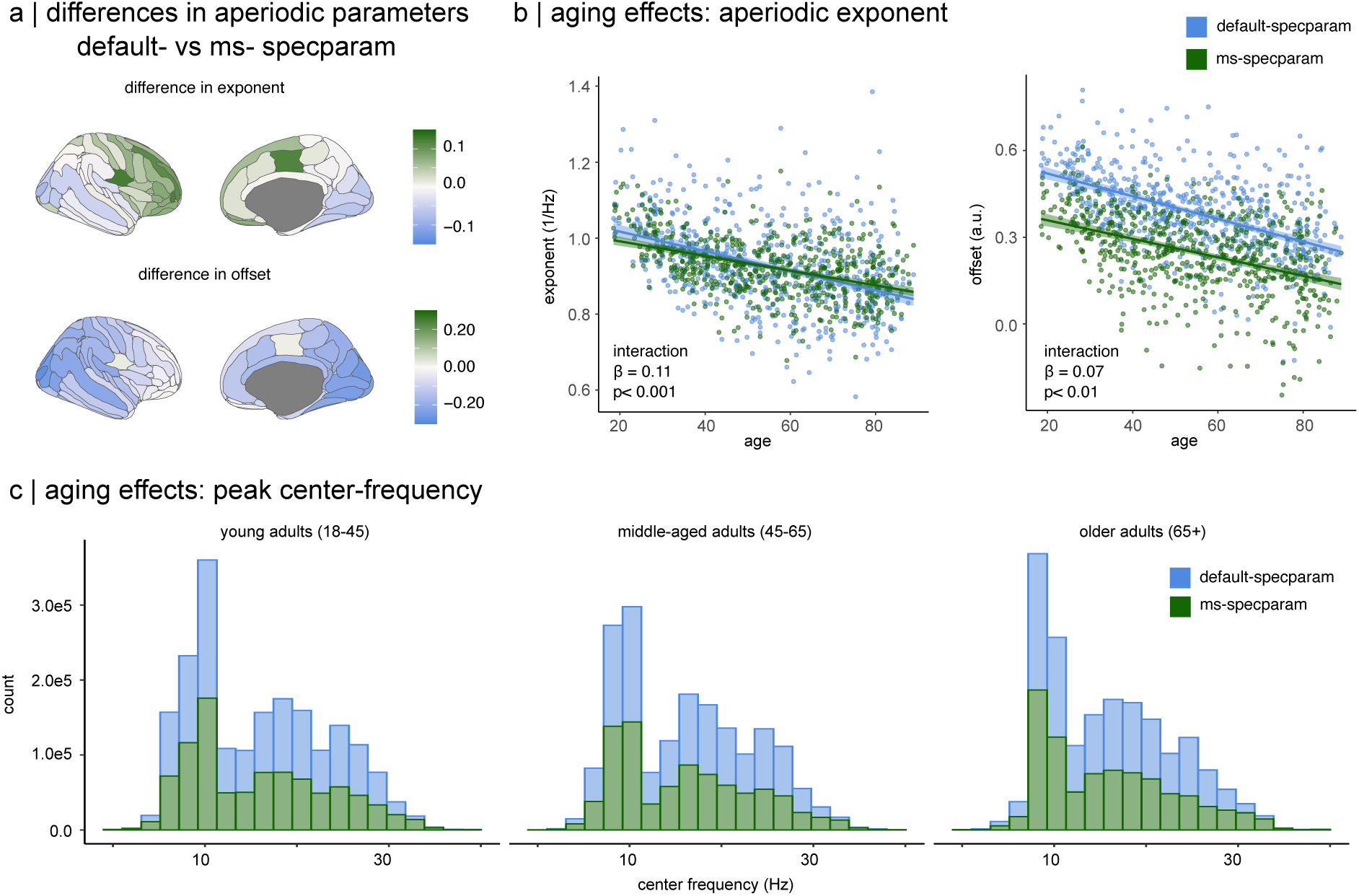
Age-related Neural Spectral Changes and Algorithmic Parsimony. (a) Topographical differences in aperiodic neural components, showing the variations in exponent (top) and offset (bottom) estimates when using *ms-specparam* versus *default-specparam*. Areas where *ms-specparam* yields higher parameter estimates are highlighted in blue. (b) Moderating effect of spectral parameterization method (*ms-specparam* vs. *Default-specparam*) on the relationship between age and aperiodic spectral components. The left graph shows the exponent, and the right graph displays the offset, with statistical interactions highlighted. (c) Frequency-specific empirical distribution of the number of peaks fitted across different age groups. The histograms show that *ms-specparam* (green) generally fits fewer peaks, especially in the mid-frequency range (8-30Hz), illustrating a more parsimonious approach to model fitting and potentially more accurate reflection of age-related spectral changes.

We obtained similar results for the age-related decline in aperiodic offset. Our analysis confirmed a significant interaction effect between age and spectral parameterization algorithm, when comparing *ms-specparam* to *default-specparam* (β = 0.07, SE = 0.03, CI [0.02, 0.13]; Figure 5b and Table S2).

Taken together, these observations suggest that the magnitude of age-related declines in the aperiodic exponent and offset are influenced by the choice of spectral parameterization algorithm. Note that the effect size of age was smaller in the spectral models derived from *ms-specparam* (exponent: β = −0.37 vs β = −0.48; offset: β = −0.40 vs β = −0.47) than *default-specparam*.

As in our simulation results, we also found that *ms-specparam* identified fewer spectral peaks than *default-specparam,* particularly in the 8-30 Hz range. This reinforces the earlier finding that *ms-specparam* curtails the number of detected peaks and underscores the influence of model selection on the characterization of peak parameters across all age groups (Figure 5c).

## 4 Discussion

Spectral parameterization enables the spectral decomposition of neurophysiological signals into aperiodic and periodic components. Its adoption has grown rapidly over recent years, thanks to open software tools, with the aim of disambiguating the respective functions of rhythmic oscillatory and arrhythmic background neural activity. However, present tools require the manual adjustment of algorithm parameters (hyperparameters), which hinders the reproducibility, interpretability, and proper fitting of spectral parameterization models to the empirical data. In the present report, we addressed this issue with the addition of a principled model selection strategy to *specparam* (*ms-specparam*) to simplify setting key hyperparameters, notably the maximum number of peaks. We validated our approach with synthetic and empirical data. We show that the resulting spectral parameterizations are more parsimonious and fit better the data, while being considerably less dependent on user decisions and expertise.

### 4.1 Spectral Parameterization with Enhanced Model Parsimony and Goodness-of-Fit

Our examination of ground-truth data shows that *ms-specparam* is more effective that *specparam* in accurately identifying spectral peaks. It avoids the frequent issue in practice of overfitting the spectral data, which typically overestimates the number of periodic components in power spectra (Figure 2a). In empirical data also, *ms-specparam* consistently generates more parsimonious models (Figure 4). We observed enhanced goodness-of-fit (reduced residual variance) in both synthetic and real-world data (Figures 2b and 4a), particularly at the edges of the frequency spectrum. The implications of these findings are twofold: firstly, *ms-specparam*’s parsimonious approach prevents both overfitting and *underfitting* (see Practical Guidelines below). Secondly, the improved accuracy of aperiodic component estimates is critical for characterizing complex neural dynamics (Donoghue et al., 2020; Gerster et al., 2022).

While we did not observe that the distance between synthetic spectral peaks influenced the ability of *ms-specparam* to detect peaks, lower amplitude peaks were on average missed more frequently than their higher amplitude counterparts for both algorithms (Figure 3c). This was particularly evident for synthetic data containing four simulated peaks. We interpret these findings as follows: as the number of peaks increases, the marginal benefit of an additional peak to the overall model fit is reduced. This is especially true because peaks are fit in decreasing order of amplitude. While both algorithms show this sensitivity bias, *ms-specparam* exhibited the lowest sensitivity for low-amplitude peaks.

In cases where data may contain multiple low-amplitude peaks, such as in infant EEG studies, users may prefer the higher positive predictive value of *ms-specparam* over the higher sensitivity of *default-specparam*, particularly when false positive peaks are of greater concern (e.g., when testing for the presence or absence of oscillatory activity).”

An interesting and not inherently obvious effect of model selection to optimize the number of peaks fit to the power spectrum is the reduced residual variance and error in parameter estimates, observed in both the synthetic and empirical data. Our interpretation of this effect is twofold.

First, *ms-specparam* simultaneously optimizes all spectral parameters—including both aperiodic and periodic—which may permit shifting some model variance between the two components that make up the power spectrum. *Second, ms-specparam* fits higher amplitude peaks with larger bandwidths in comparison to *default-specparam*. As observed in the example spectrum plotted in Figure 3B, this yields more parsimonious model fits. We speculate that by removing these narrow-band low amplitude peaks fit by *default-specparam*, *ms-specparam* reassigns this part of this variance to improve the final fit of the aperiodic component of the neural power spectrum. This results, therefore in better overall fits and a reduction in model fit error, principally along the edges of the power spectrum (see Figure 3A).

Note that although *ms-specparam* fits models with less residual variance that are more parsimonious than *default-specparam*, the model selection algorithm takes approximately 8 times longer to run on the empirical data. *Default-specparam*, on average, fit the 148 spectra of a participant in 3.01 seconds (s.d. = 0.48), whereas *ms-specparam* took 24.51 seconds (s.d. = 5.95). These benchmarks were obtained on a standard desktop machine. While *ms-specparam* is therefore slower, the cost remains practical for most research applications, and can be reduced through parallelization or the use of high-performance computing infrastructure.

### 4.2 Hyperparameter Settings

Our empirical investigations emphasize the critical role of hyperparameter settings in spectral modelling. Notably, the choice of spectral parameterization algorithm influenced aperiodic parameter estimates across methods (Figures 5a-c). While deviations were modest, suggesting that default spectral parameterizations may have captured the aperiodic component accurately, our comparative analysis with synthetic data indicates that *ms-specparam* yields improved estimates of the aperiodic exponent and offset (Figure 2b).

We replicated with the new model selection approach previous observations of changing aperiodic parameters with age (Cellier et al., 2021; Donoghue et al., 2020; Hill et al., 2022). We found that the aperiodic component of the neurophysiological power spectrum flattens with age, which has been discussed as related to increased neural noise and asynchronous neuronal firing, yielding less structured brain dynamics (Usher et al., 1995; Pozzorini et al., 2013; Voytek et al., 2015; Voytek & Knight, 2015). Several empirical observations support this hypothesis (Bédard et al., 2006; Sosnoff & Newell, 2011).

We found, however, that previously reported effects of shifts in the spectral aperiodic exponent and offset with age are reduced, depending on the spectral parameterization method used and its hyperparameters. This observation encourages the present and further efforts towards more automated and principled parameter selection procedures, promoting robustness and replicability of research results.

### 4.3 Practical Guidelines

We provide practical recommendations to adjust the hyperparameters of *ms-specparam*. As with other spectral parameterization methods, we encourage future users to examine their data’s power spectra before and after applying *ms-specparam* and verify the model’s goodness-of-fit. We refer the reader to the previously published guidelines by Donoghue et al. (2020), Gerster et al. (2022), and Ostlund et al. (2022), which set good foundational principles for neurophysiological spectral parameterization. Here, we highlight more specific considerations for the best possible use of *ms-specparam*:

1. Model selection in *ms-specparam* determines the optimal number of spectral peaks that fit the empirical power spectrum. The settings of other hyperparameters, including peak width limits and aperiodic mode, remain to be defined by the user, but should always reflect the qualities of the data upon visual inspection. The value set for the maximum number of spectral peaks parameter needs to be larger than the number of peaks that are clearly visible in the data’s power spectrum. In our investigations, we set this value to 6.
2. We encourage users to derive measures of model goodness-of-fit, such as Mean-Squared Error, R^2^, and BIC, as in our *Brainstorm* plug-in of the proposed model selection methods. Some of these metrics, like R^2^, may be influenced by the aperiodic component of the spectrum, as demonstrated in previous studies (Donoghue et al., 2020). In the context of model selection, users should choose the spectral model with the lowest BIC (as in the present study).

To conclude, the present report introduced a model selection approach to the parameterization of neurophysiological power spectra. The approach minimizes the requirement for user expertise in the adjustment of hyperparameters, which influences the outcome of analyses. It is grounded in the optimization of parameters that favour model parsimony while maximizing goodness-of-fit. We foresee that this principled approach will contribute to the robust application of spectral parameterization in neuroscience research, further elucidating the roles of rhythmic and arrhythmic brain activity in cognition, health, and disease. We anticipate that *ms-specparam* will enhance the reproducibility and robustness of reported findings.

## Data and Software Availability

The *ms-specparam* algorithm, as well as simulated power spectra and code used to generate results, are available on GitHub (github.com/lucwilson/model_selection). Resting-state MEG recordings were obtained from the CamCAN repository (Shafto et al., 2014; Taylor et al., 2017).

## Author Contributions

Conceptualization: LEW, JDSC

Data Curation: LEW, JDSC

Methodology: LEW, JDSC, BLK

Software: LEW, JDSC

Visualization: LEW, JDSC, BLK

Funding acquisition: SB

Writing – original draft: LEW, JDSC, SB

Writing – review & editing: LEW, JDSC, BLK, SB

## Declaration of Competing Interests

All authors declare no competing conflicts of interest. The listed funding sources in the **Acknowledgements** did not play any role in the writing of the manuscript or the decision to submit this manuscript for publication.

## Acknowledgements

SB is supported by the NIH (R01EB026299-05), the Tier-1 CIHR Canada Research Chair of Neural Dynamics of Brain Systems (CRC-2017-00311), a Discovery Grant from the Natural Sciences and Engineering Research Council of Canada (436355-13), and the Canadian Institutes of Health Research (202503-PJT-542788). This work was also supported by a doctoral fellowship from NSERC (JDSC), as well as by a master’s fellowship from NSERC (LEW).

## Supplementary Materials

### Algorithm Validation with Alternative Noise Conditions

In addition to injecting moderate noise to the synthetic data, we also evaluated *ms-specparam* performance at low (s.d. = 0.02) and high (s.d. = 0.10) noise conditions. In spectra with low noise, *ms-specparam* detected spectral peaks with comparable sensitivity (92%) to *default-specparam* (92%). The positive predictive value of peaks detected using *ms-specparam* (97%) was notably higher than that of peaks detected using *default-specparam* (68%). In spectra with high noise, *ms-specparam* detected spectral peaks with lower sensitivity (84%) to *default-specparam* (90%). However, the positive predictive value of peaks detected using *ms-specparam* (96%) remained higher than that of peaks detected using *default-specparam* (61%). Taken together, these results support the generalizability of the observed algorithmic performance improvements across noise levels.

**Figure S1:**
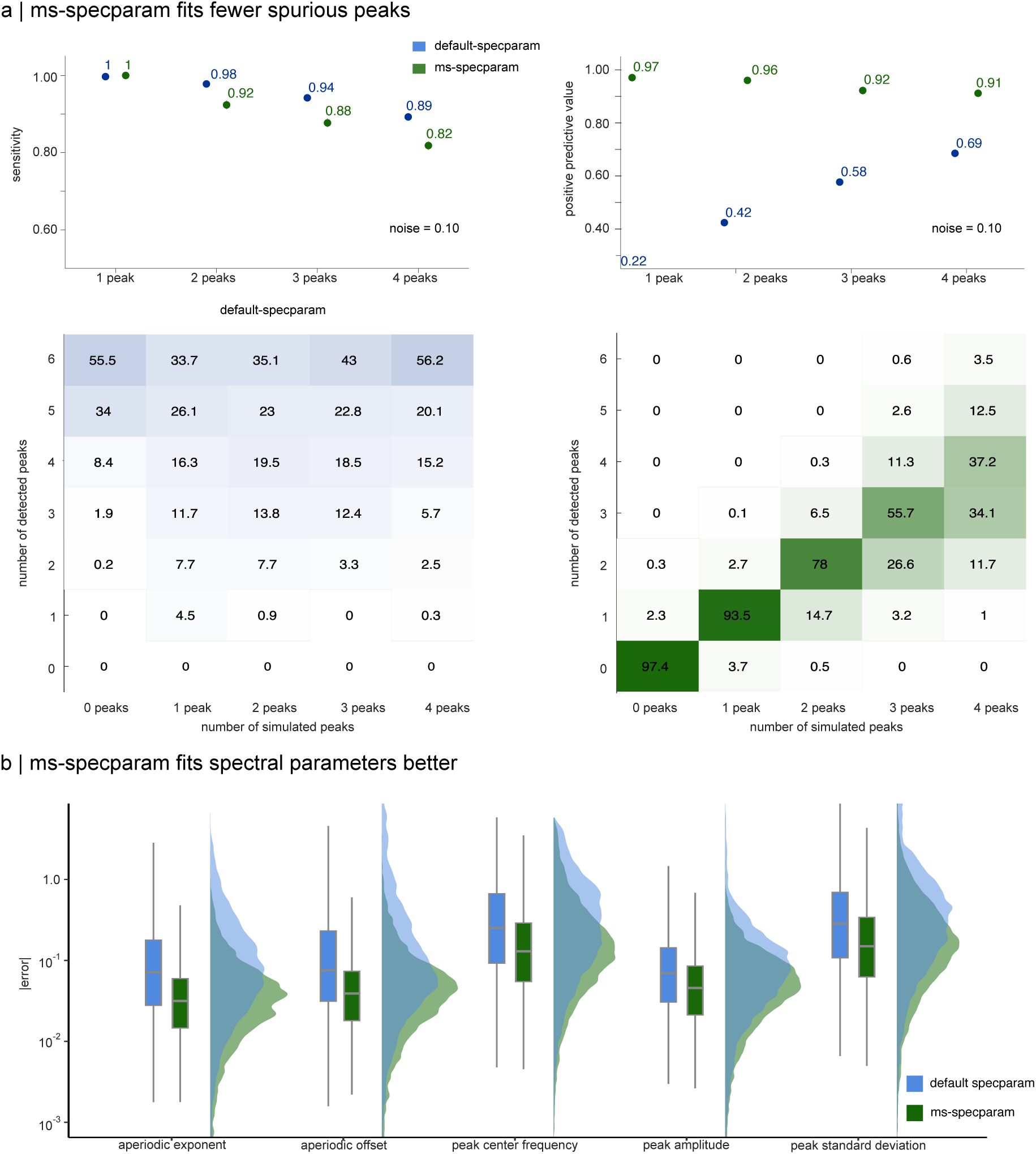
Performance of *ms-specparam* and *default-specparam* on synthetic stationary data at higher noise levels (SNR = 0.10). (a) Sensitivity and positive predictive value (PPV) for spectral peak detection across simulations with high noise. While *ms-specparam* (green) shows slightly lower sensitivity than *default-specparam* (blue), it consistently achieves higher PPV, indicating greater reliability in detected peaks. The accompanying heatmaps show the correspondence between the ground-truth and estimated number of peaks (values indicate percentage incidence), illustrating *ms-specparam*’s improved accuracy in peak count estimation. (b) Boxplots and empirical density plots of absolute estimation errors for spectral parameters (aperiodic exponent and offset, peak center frequency, amplitude, and bandwidth). *ms-specparam* yields lower estimation errors across all parameters compared to *default-specparam*.

**Figure S2:**
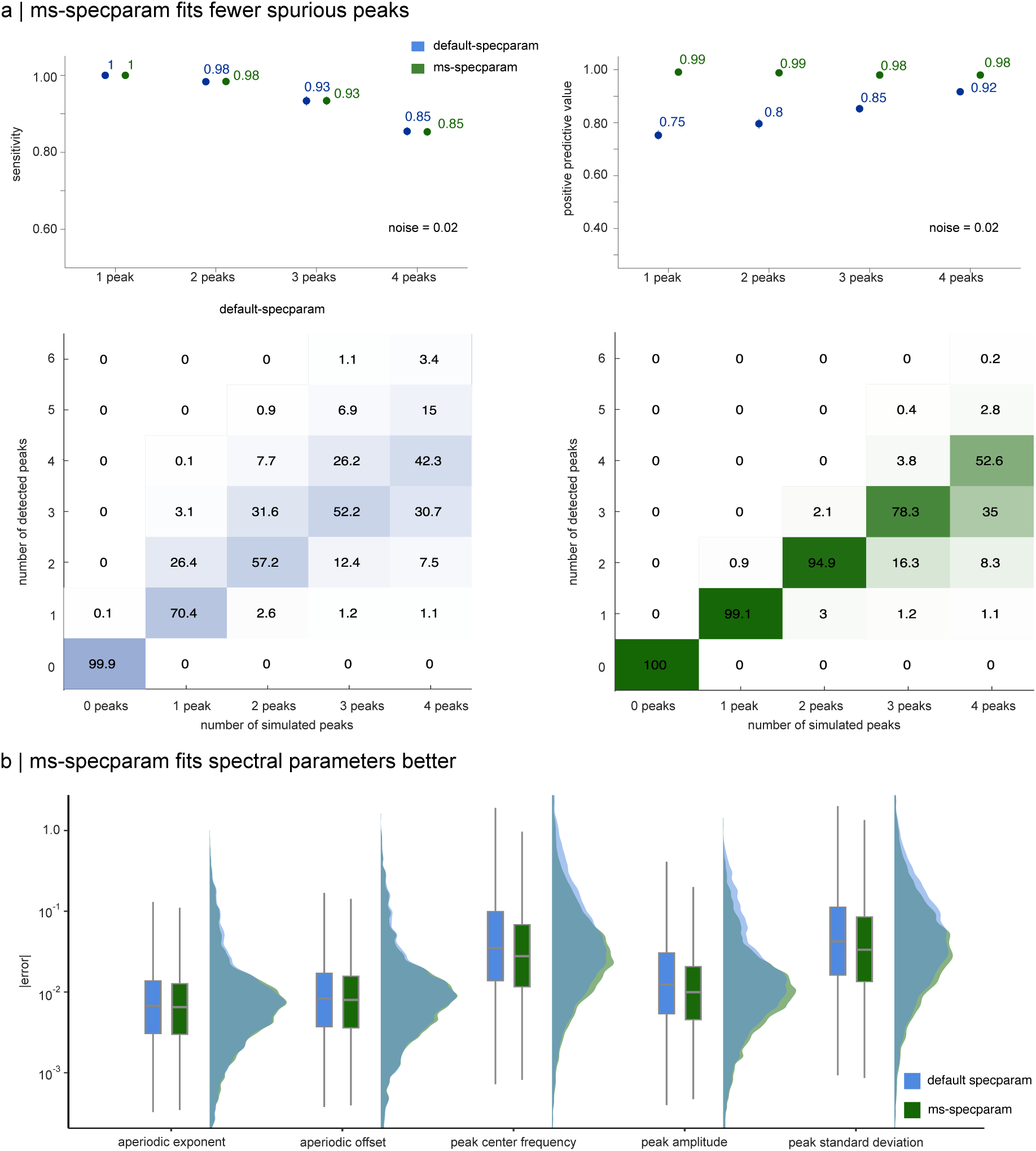
Performance of *ms-specparam* and *default-specparam* on synthetic stationary data at lower noise levels (SNR = 0.02). (a) Sensitivity and positive predictive value (PPV) for spectral peak detection across simulations with low noise. *ms-specparam* (green) and *default-specparam* (blue) show comparable sensitivity, but *ms-specparam* consistently achieves higher PPV, indicating fewer false positives. The accompanying heatmaps display the correspondence between ground-truth and estimated number of peaks (values indicate percentage incidence), highlighting *ms-specparam*’s improved precision in estimating peak count. (b) Boxplots and empirical density plots of absolute estimation errors for spectral parameters (aperiodic exponent and offset, peak center frequency, amplitude, and bandwidth). *ms-specparam* outperforms *default-specparam* by producing lower estimation errors across all parameter types.

### Empirical MEG Data

**Figure S3:**
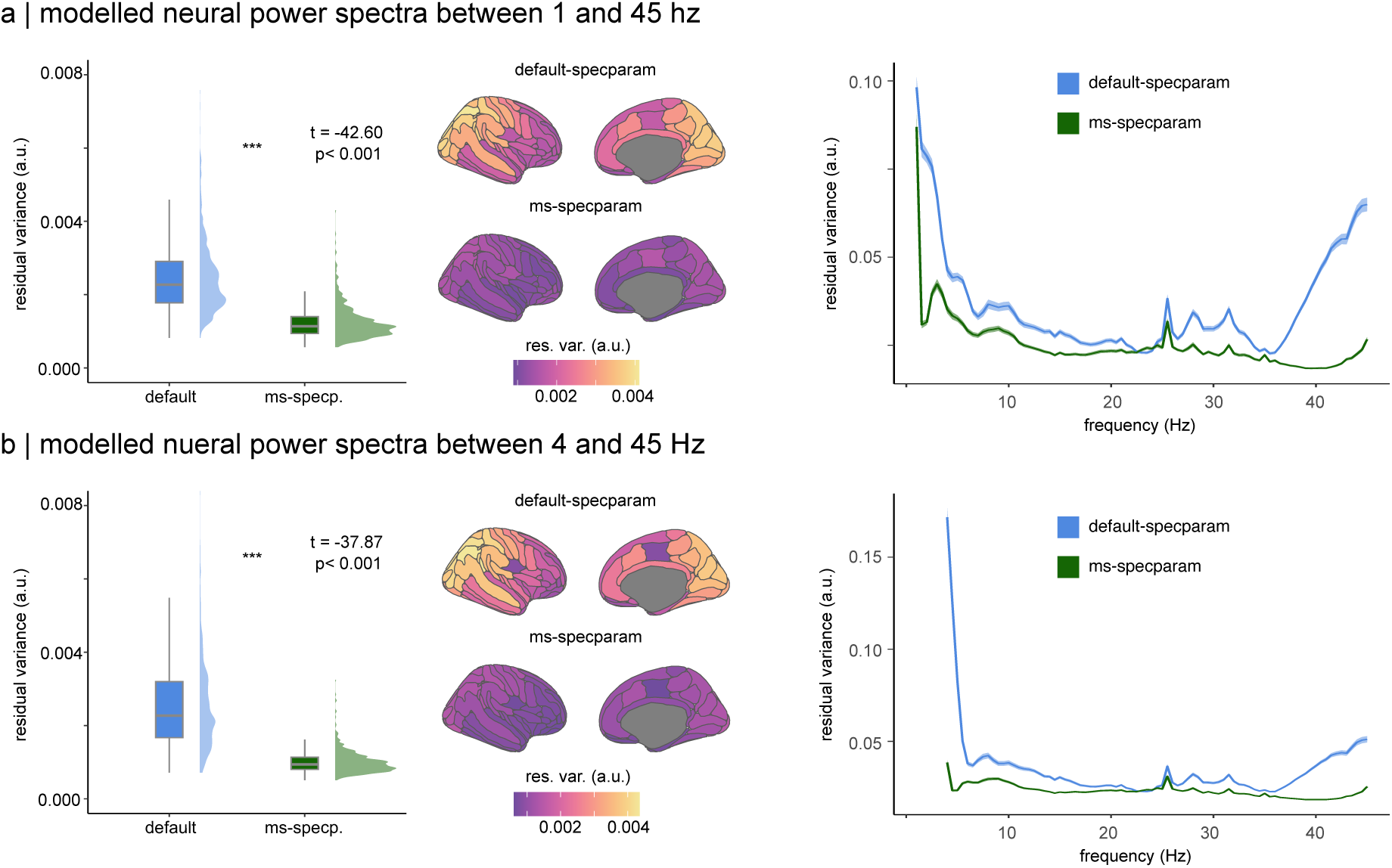
*ms-specparam* is Robust Against Choice of Frequency Ranges to be Fit. We tested the robustness of *ms-specparam* against the choice of the frequency range of the spectrum to fit. Neural power spectra were parameterized between 1 and 45Hz (a) or 4 and 45 Hz (b) using both algorithms. *ms-specparam* (green) consistently achieved lower residual variance, indicating a superior fit, relative to *default*-*specparam* (blue; boxplot) regardless of the frequency range parameterized. The brain maps display residual variance values for each cortical parcel. A frequency breakdown (right panel) reveals *ms-specparam* outperforms default *specparam* variations across the spectrum, particularly at the edges of the frequency spectrum.

**Table S1.**
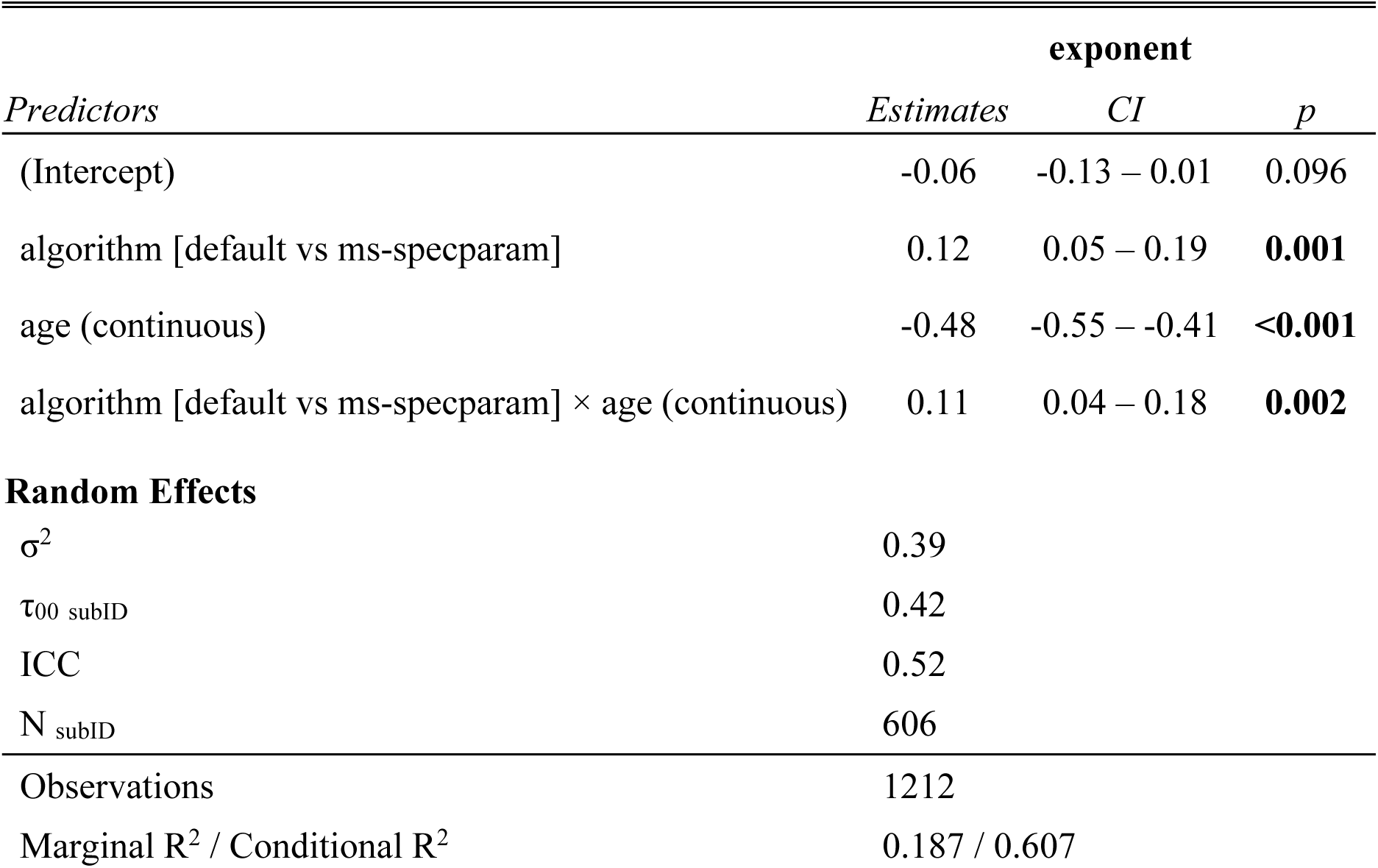
Hyperparameter choice impacts the age’s effect on the aperiodic exponent: default hyperparameter setting vs *ms-specparam*.

| *Predictors* | **exponent** |  |  |
| --- | --- | --- | --- |
|  | *Estimates* | *CI* | *p* |
| (Intercept) | -0.06 | -0.13 – 0.01 | 0.096 |
| algorithm [default vs ms-specparam] | 0.12 | 0.05 – 0.19 | **0.001** |
| age (continuous) | -0.48 | -0.55 – -0.41 | **<0.001** |
| algorithm [default vs ms-specparam] × age (continuous) | 0.11 | 0.04 – 0.18 | **0.002** |
| **Random Effects** |  |  |  |
| $\sigma^2$ | 0.39 | | |
| $\tau_{00 \text{ subID}}$ | 0.42 | | |
| ICC | 0.52 |  |  |
| $N_{\text{subID}}$ | 606 | | |
| Observations | 1212 |  |  |
| Marginal $R^2$ / Conditional $R^2$ | 0.187 / 0.607 | | |

**Table S2.**
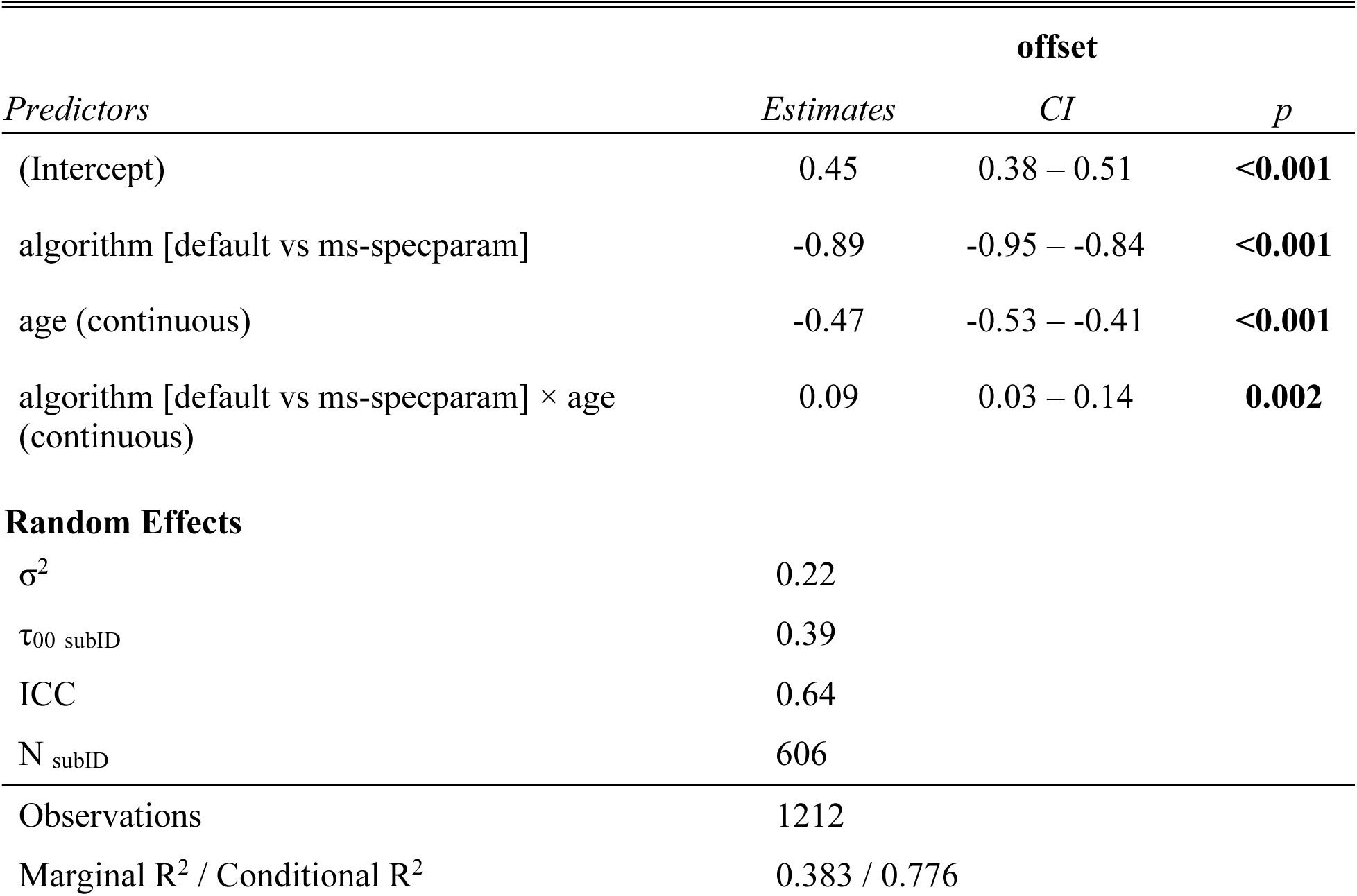
Hyperparameter choice impacts the age’s effect on the aperiodic offset: default hyperparameter setting vs *ms-specparam*.

## Notes

https://www.github.com/lucwilson/model_selection

### Competing Interest Statement

The authors have declared no competing interest.

### Summary of Updates

Modifications to main text, substitutions of figures, and improvements to article clarity.

## References

Bédard, C., Kröger, H., & Destexhe, A. (2006). Does the 1/f frequency scaling of brain signals reflect self-organized critical states? Physical Review Letters, 97(11), 118102. doi:10.1103/PhysRevLett.97.118102

Brady, B., & Bardouille, T. (2022). Periodic/aperiodic parameterization of transient oscillations (PAPTO)-Implications for healthy ageing. NeuroImage, 251, 118974. doi:10.1016/j.neuroimage.2022.118974

Brake, N., Duc, F., Rokos, A., Arseneau, F., Shahiri, S., Khadra, A., & Plourde, G. (2024). A neurophysiological basis for aperiodic EEG and the background spectral trend. Nature Communications, 15(1), 1514. doi:10.1038/s41467-024-45922-8

Buzsáki, G., & Watson, B. O. (2012). Brain rhythms and neural syntax: Implications for efficient coding of cognitive content and neuropsychiatric disease. Dialogues in Clinical Neuroscience, 14(4), 345–367. doi:10.31887/DCNS.2012.14.4/gbuzsaki

Cellier, D., Riddle, J., Petersen, I., & Hwang, K. (2021). The development of theta and alpha neural oscillations from ages 3 to 24 years. Developmental Cognitive Neuroscience, 50, 100969. doi:10.1016/j.dcn.2021.100969

Chini, M., Pfeffer, T., & Hanganu-Opatz, I. (2022). An increase of inhibition drives the developmental decorrelation of neural activity. eLife, 11, e78811. doi:10.7554/eLife.78811

da Silva Castanheira, J., Wiesman, A. I., Hansen, J. Y., Misic, B., Baillet, S., Breitner, J.,…Vitali, P. (2024). The neurophysiological brain-fingerprint of Parkinson’s disease. eBioMedicine, 105. doi:10.1016/j.ebiom.2024.105201

da Silva Castanheira, J., Wiesman, A. I., Taylor, M. J., & Baillet, S. (2024). The Lifespan Evolution of Individualized Neurophysiological Traits. bioRxiv, 2024.2011.2027.624077. doi:10.1101/2024.11.27.624077

Destrieux, C., Fischl, B., Dale, A., & Halgren, E. (2010). Automatic parcellation of human cortical gyri and sulci using standard anatomical nomenclature. NeuroImage, 53(1), 1–15. doi:10.1016/j.neuroimage.2010.06.010

Donoghue, T., Haller, M., Peterson, E. J., Varma, P., Sebastian, P., Gao, R.,…Voytek, B. (2020). Parameterizing neural power spectra into periodic and aperiodic components. Nature Neuroscience, 23(12), 1655–1665. doi:10.1038/s41593-020-00744-x

Fischl, B. (2012). FreeSurfer. NeuroImage, 62(2), 774–781. doi:10.1016/j.neuroimage.2012.01.021

Gao, R., Peterson, E. J., & Voytek, B. (2017). Inferring synaptic excitation/inhibition balance from field potentials. NeuroImage, 158, 70–78. doi:10.1016/j.neuroimage.2017.06.078

Gerster, M., Waterstraat, G., Litvak, V., Lehnertz, K., Schnitzler, A., Florin, E.,…Nikulin, V. (2022). Separating neural oscillations from aperiodic 1/f activity: Challenges and recommendations. Neuroinformatics. doi:10.1007/s12021-022-09581-8

Gross, J., Baillet, S., Barnes, G. R., Henson, R. N., Hillebrand, A., Jensen, O.,…Schoffelen, J.-M. (2013). Good practice for conducting and reporting MEG research. NeuroImage, 65, 349–363. doi:10.1016/j.neuroimage.2012.10.001

Gyurkovics, M., Clements, G. M., Low, K. A., Fabiani, M., & Gratton, G. (2022). Stimulus-induced changes in 1/f-like background activity in EEG. The Journal of Neuroscience, 42(37), 7144. doi:10.1523/JNEUROSCI.0414-22.2022

Hill, A. T., Clark, G. M., Bigelow, F. J., Lum, J. A. G., & Enticott, P. G. (2022). Periodic and aperiodic neural activity displays age-dependent changes across early-to-middle childhood. Developmental Cognitive Neuroscience, 54, 101076. 10.1016/j.dcn.2022.101076

Jefferys, W. H., & Berger, J. O. (1992). Ockham’s Razor and Bayesian Analysis. American Scientist, 80(1), 64–72. Retrieved from http://www.jstor.org/stable/29774559.

Kosciessa, J. Q., Grandy, T. H., Garrett, D. D., & Werkle-Bergner, M. (2020). Single-trial characterization of neural rhythms: Potential and challenges. NeuroImage, 206, 116331. doi:10.1016/j.neuroimage.2019.116331

Miller, K. J., Honey, C. J., Hermes, D., Rao, R. P. N., denNijs, M., & Ojemann, J. G. (2014). Broadband changes in the cortical surface potential track activation of functionally diverse neuronal populations. NeuroImage, 85, 711–720. doi:10.1016/j.neuroimage.2013.08.070

Mitchell, T. M. (1997). Machine Learning. New York: McGraw-Hill.

Myung, I. J. (2000). The importance of complexity in model selection. Journal of Mathematical Psychology, 44(1), 190–204. doi:10.1006/jmps.1999.1283

Ostlund, B., Donoghue, T., Anaya, B., Gunther, K. E., Karalunas, S. L., Voytek, B., & Pérez-Edgar, K. E. (2022). Spectral parameterization for studying neurodevelopment: How and why. Developmental Cognitive Neuroscience, 54, 101073. doi:10.1016/j.dcn.2022.101073

Ostlund, B. D., Alperin, B. R., Drew, T., & Karalunas, S. L. (2021). Behavioral and cognitive correlates of the aperiodic (1/f-like) exponent of the EEG power spectrum in adolescents with and without ADHD. Developmental Cognitive Neuroscience, 48, 100931. doi:10.1016/j.dcn.2021.100931

Pozzorini, C., Naud, R., Mensi, S., & Gerstner, W. (2013). Temporal whitening by power-law adaptation in neocortical neurons. Nature Neuroscience, 16(7), 942–948. doi:10.1038/nn.3431

Preston, M., Schaworonkow, N., & Voytek, B. (2022). Oscillations and aperiodic activity: Evidence for dynamic changes in both during memory encoding. bioRxiv, 2022.2010.2004.509632. doi:10.1101/2022.10.04.509632

Schwarz, G. (1978). Estimating the dimension of a model. The Annals of Statistics, 6(2), 461–464. doi:10.1214/aos/1176344136

Seymour, R. A., Alexander, N., & Maguire, E. A. (2022). Robust estimation of 1/f activity improves oscillatory burst detection. European Journal of Neuroscience, 56(10), 5836–5852. doi:10.1111/ejn.15829

Shafto, M. A., Tyler, L. K., Dixon, M., Taylor, J. R., Rowe, J. B., Cusack, R.,…Cam-CAN (2014). The Cambridge Centre for Ageing and Neuroscience (Cam-CAN) study protocol: A cross-sectional, lifespan, multidisciplinary examination of healthy cognitive ageing. BMC Neurology, 14(1), 204. doi:10.1186/s12883-014-0204-1

Sosnoff, J. J., & Newell, K. M. (2011). Aging and motor variability: A test of the neural noise hypothesis. Experimental Aging Research, 37(4), 377–397. doi:10.1080/0361073X.2011.590754

Stokes, P. A., Rath, P., Possidente, T., He, M., Purcell, S., Manoach, D. S.,…Prerau, M. J. (2023). Transient oscillation dynamics during sleep provide a robust basis for electroencephalographic phenotyping and biomarker identification. Sleep, 46(1), zsac223. doi:10.1093/sleep/zsac223

Tadel, F., Baillet, S., Mosher, J. C., Pantazis, D., & Leahy, R. M. (2011). Brainstorm: A user-friendly application for MEG/EEG analysis. Computational Intelligence and Neuroscience, 2011, 879716. doi:10.1155/2011/879716

Taylor, J. R., Williams, N., Cusack, R., Auer, T., Shafto, M. A., Dixon, M.,…Henson, R. N. (2017). The Cambridge Centre for Ageing and Neuroscience (Cam-CAN) data repository: Structural and functional MRI, MEG, and cognitive data from a cross-sectional adult lifespan sample. NeuroImage, 144, 262–269. doi:10.1016/j.neuroimage.2015.09.018

Usher, M., Stemmler, M., & Olami, Z. (1995). Dynamic pattern formation leads to 1/f noise in neural populations. Physical Review Letters, 74(2), 326–329. doi:10.1103/PhysRevLett.74.326

Vandekerckhove, J., Matzke, D., & Wagenmakers, E.-J. (2015). Model comparison and the principle of parsimony. In The Oxford Handbook of Computational and Mathematical Psychology. doi:10.1093/oxfordhb/9780199957996.013.14

Voytek, B., & Knight, R. T. (2015). Dynamic network communication as a unifying neural basis for cognition, development, aging, and disease. Biological Psychiatry, 77(12), 1089–1097. doi:10.1016/j.biopsych.2015.04.016

Voytek, B., Kramer, M. A., Case, J., Lepage, K. Q., Tempesta, Z. R., Knight, R. T., & Gazzaley, A. (2015). Age-related changes in 1/f neural electrophysiological noise. The Journal of Neuroscience, 35(38), 13257. doi:10.1523/JNEUROSCI.2332-14.2015

Wagenmakers, E.-J. (2007). A practical solution to the pervasive problems of p values. Psychonomic Bulletin & Review, 14(5), 779–804. doi:10.3758/BF03194105

Waschke, L., Donoghue, T., Fiedler, L., Smith, S., Garrett, D. D., Voytek, B., & Obleser, J. (2021). Modality-specific tracking of attention and sensory statistics in the human electrophysiological spectral exponent. eLife, 10, e70068. doi:10.7554/eLife.70068

Wen, H., & Liu, Z. (2016). Separating fractal and oscillatory components in the power spectrum of neurophysiological signal. Brain Topography, 29(1), 13–26.

Whitten, T. A., Hughes, A. M., Dickson, C. T., & Caplan, J. B. (2011). A better oscillation detection method robustly extracts EEG rhythms across brain state changes: The human alpha rhythm as a test case. NeuroImage, 54(2), 860–874. doi:10.1016/j.neuroimage.2010.08.064

Wiesman, A. I., da Silva Castanheira, J., & Baillet, S. (2022). Stability of spectral estimates in resting-state magnetoencephalography: Recommendations for minimal data duration with neuroanatomical specificity. NeuroImage, 247, 118823. doi:10.1016/j.neuroimage.2021.118823

Wiest, C., Torrecillos, F., Pogosyan, A., Bange, M., Muthuraman, M., Groppa, S.,…Tan, H. (2023). The aperiodic exponent of subthalamic field potentials reflects excitation/inhibition balance in Parkinsonism. eLife, 12, e82467. doi:10.7554/eLife.82467

Wilson, L. E., da Silva Castanheira, J., & Baillet, S. (2022). Time-resolved parameterization of aperiodic and periodic brain activity. eLife, 11, e77348. doi:10.7554/eLife.77348

